# Resurrected nitrogenases recapitulate canonical N-isotope biosignatures over two billion years

**DOI:** 10.1101/2025.05.21.655380

**Authors:** Holly Rucker, Kunmanee Bubphamanee, Derek F. Harris, Kurt Konhauser, Lance C. Seefeldt, Roger Buick, Betül Kaçar

## Abstract

Nitrogen isotope fractionation (ε^15^N) in sedimentary rocks has provided evidence for biological nitrogen fixation, and thus primary productivity, on the early Earth. However, the extent to which molecular evolution has influenced the isotopic signatures of nitrogenase, the enzyme that catalyzes the conversion of atmospheric nitrogen (N_2_) to bioavailable ammonia, remains unresolved. Here, we reconstruct and experimentally characterize a library of synthetic ancestral nitrogenase genes, spanning over 2 billion years of evolutionary history. By engineering modern microbes to express these ancient nitrogenases, we assess the resulting ε^15^N values under controlled laboratory conditions. All engineered strains exhibit ε^15^N values within a narrow range comparable to that of modern microbes, suggesting that molybdenum (Mo)-dependent nitrogenase has been largely invariant throughout evolutionary time since the origins of this pathway. These results confirm the robustness of N-isotope biosignatures in the ancient rock record and bolster their utility in the search for life in extraterrestrial environments.

## INTRODUCTION

The metabolic and physiological characteristics of Earth’s earliest life remain largely obscure, with only the broad outlines discernible for the first 2 billion years of evolution^1–7^. In the absence of diagnostic fossils, stable isotopes of carbon and sulfur have served as key geochemical proxies for ancient biological activity^2,8–11^. Nitrogen, owing to its centrality in biochemistry, is an appealing candidate for early life reconstructions. However, the complexity and diversity of its metabolic pathways have hindered the interpretation of its record. Uniquely, nitrogen fixation of N_2_ into bioavailable N is catalyzed by a single enzyme, nitrogenase. This constraint raises the possibility that the evolutionary history of this single enzyme could inform interpretation of nitrogen isotopes across deep time.

The oldest geochemical evidence for biological nitrogen fixation comes from nitrogen isotopic compositions of ∼3.2 Ga Archean sedimentary rocks, with δ^15^N values ranging from +0.7‰ to - 2.8‰^12^. These values closely resemble those of the biomass of modern organisms using Mo-dependent nitrogenase, which typically fall between +1‰ and −3‰^12–16^. By contrast, modern organisms expressing alternative vanadium (V)- and iron (Fe)-based nitrogenase isozymes exhibit more negative biomass fractionations (−6‰ to −8‰)^13^. Similar isotopic patterns in ∼2.9 Ga rocks have been interpreted as evidence for the primordial nitrogen cycle driven by Mo-nitrogenase^17^. Additionally, nitrogen isotope signatures from transient atmospheric oxygenation episodes (“whiffs”) at ∼2.66 Ga and ∼2.5 Ga suggest that Mo-nitrogenase remained the dominant nitrogen-fixing enzyme before the onset of the Great Oxidation Event (GOE)^9,20^ at approximately 2.45 Ga, after which time the atmosphere became permanently oxygenated^18^. Although post-GOE oxidative processes increasingly overprint nitrogen fixation signals in younger rocks, Mesoproterozoic offshore marine sediments deposited under low-oxygen conditions still preserve δ^15^N values consistent with Mo-nitrogenase activity^19–21^. Even during Phanerozoic ocean anoxic events, such as those in the late Devonian, Jurassic, and Cretaceous, marine sedimentary δ^15^N values often return to the canonical Mo-nitrogenase range^22^. Collectively, these records depend on a critical assumption: that nitrogenase-mediated fractionation has remained stable over billions of years, despite profound environmental, ecological, and evolutionary shifts.

Isotopic fractionation during biological nitrogen fixation varies across modern diazotrophs depending on species^16^, nitrogenase isozyme^13^, and environmental conditions^14,16,23,24^. Importantly, all extant nitrogenases are the products of billions of years of molecular evolution, and adapted to an Earth with an oxygenated atmosphere and an ecologically diverse biosphere^25^. These factors can impact the isotopic fractionation effect generated from nitrogenase activity by modifying the mechanistic aspects of catalysis. Therefore, it remains uncertain whether the isotopic fractionation signatures observed in modern biology can be reliably extrapolated to interpret nitrogen isotope signatures in the ancient sedimentary record.

We constructed a library of ancient nitrogenases, spanning over 2 billion years, and engineered a model diazotroph, *Azotobacter vinelandii (A. vinelandii)*, to express these ancient variants *in vivo*. Our work separates itself from prior work, as we start from synthetic DNA rather than modifying existing genes. To date, no experimental study has pursued basic genetic manipulations such as homologous gene replacements in the context of nitrogenase isotopes. In contrast, here we employ a synthetic biology approach, reconstructing nitrogenase from first principles, establishing an entirely new molecular framework to probe ancient biosignatures. We measured the nitrogen isotope fractionation in the cell biomass of engineered *A. vinelandii* strains, with all fixed nitrogen coming from the activity of the synthetic ancient nitrogenase. This system enabled a direct test of the evolutionary stability of nitrogen isotope fractionation across deep time, using controlled expression of biochemically functional, phylogenetically inferred ancient enzymes. The resulting isotopic signatures were then evaluated within the context of the sedimentary nitrogen isotopic record and long-term changes in Earth’s biogeochemistry.

## RESULTS

### Reconstruction of nitrogenase ancestors

Our study builds on a nitrogenase phylogeny and an ancestral nitrogenase protein sequence database (**Figure 1, Methods**). The phylogeny broadly samples known nitrogenase sequences and host taxonomic diversity, grouping homologs into three major clades: Groups I to III (Figure 1a). The Nif-III clade also contains two subclades corresponding to the alternative nitrogenases, Vnf and Anf, which incorporate VFe- and FeFe-cofactors, respectively, instead of the canonical FeMo-cofactor found in most nitrogenases^26^. This overall phylogenetic topology is largely consistent with prior reconstructions of nitrogenase evolution^27–29^.

**Figure 1.**
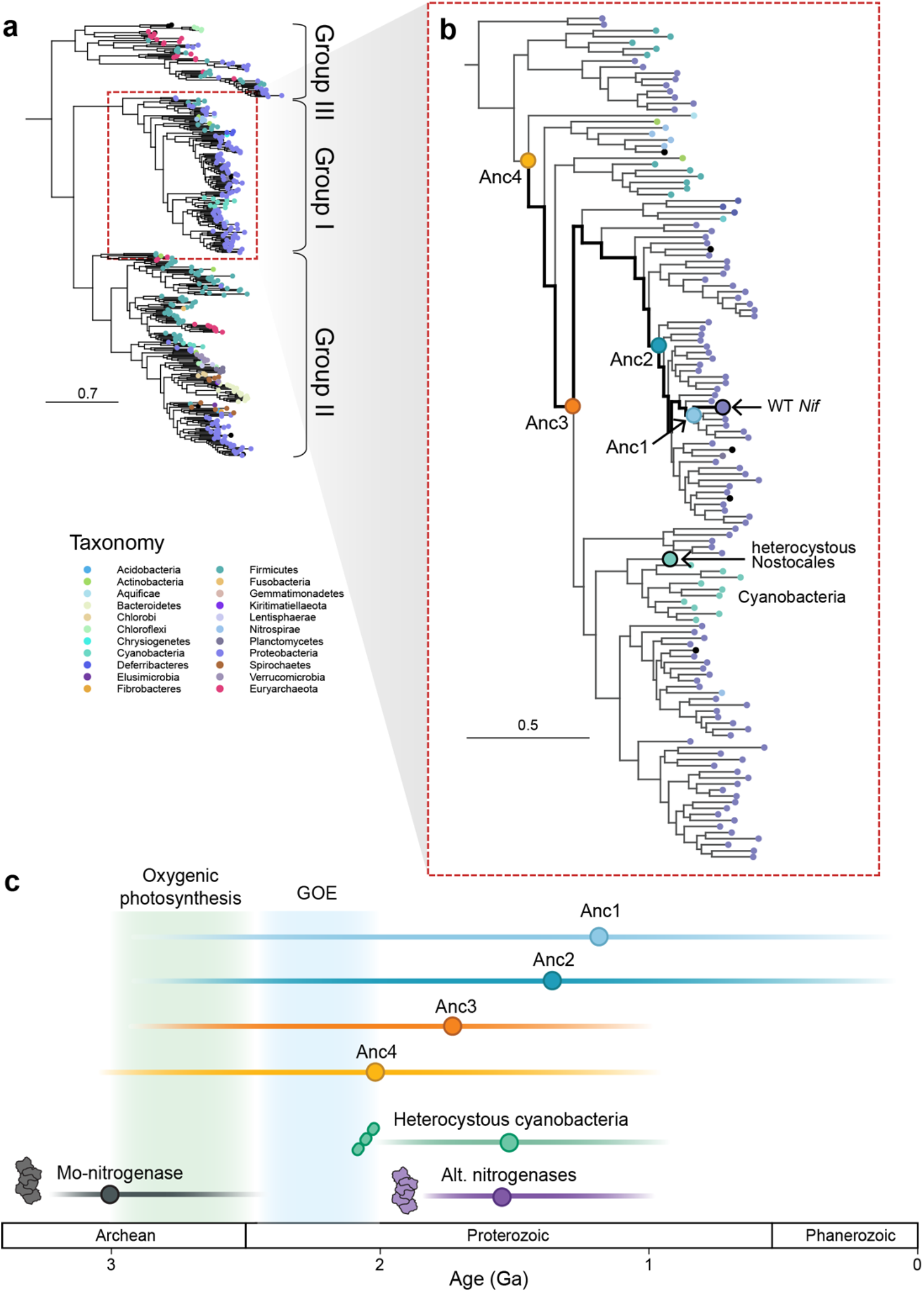
**a**, Protein phylogeny of resurrected ancestral nitrogenases (nifHDK). Node colors represent extant taxonomy. **b**, Close-up of Group I nitrogenases. The ancestral nodes targeted in this study are labeled with circles. Heterocystous cyanobacteria (highlighted in teal) serve as a minimum age constraint for the ancestral nitrogenase nodes based on fossil evidence^32^. **c**, Approximate ages for the origin of nitrogenase isoforms are based on geochemical^12^ and phylogenetic data^29^.

The Group-I clade is distinguished by its association with oxygen-tolerant hosts. In our dataset, approximately 62.5% of Nif-I nitrogenases are either found in, or predicted to occur in, aerobic or facultatively anaerobic prokaryotes, compared to only 6.5% in Group-II and 0% in Group-III (see **Methods**). This strong association between most Group-I nitrogenases and host oxygen tolerance suggests that the diversification of major Group-I lineages likely followed the initial rise of atmospheric oxygen (the GOE) and the resulting emergence of aerobic metabolism^5,30^. Notably, the Nif-I clade also contains a monophyletic cluster of nitrogenases hosted by cyanobacteria, consistent with abundant cyanobacterial microfossils preserved in Proterozoic strata^31,32^. Both observations make the Group-I clade particularly well suited for correlating nitrogenase evolution with the sedimentary rock record.

To experimentally reconstruct ancient nitrogenase isotopic signatures, we engineered four ancient nitrogenases representing a temporal series from the present into the deep past (∼700 Ma to 2.3 Ga) in a modern *A. vinelandii* strain (**Table S1**). These variants (Anc1 to Anc4), reconstructed from phylogenetic nodes along the oxygen-tolerant Group-I lineage, were individually introduced into a nitrogenase-deficient background to replace one or more native *nif* genes (**Table S1**). This inferred evolutionary context places the reconstructed ancestors after the GOE and captures a period of permanent but varying atmospheric oxygenation.

Notably, Anc3 and Anc4 precede the diversification of heterocystous cyanobacteria, which form specialized nitrogen-fixing cells^33^. Phylogenetic analysis places Anc3 prior to the emergence of a monophyletic clade that includes nitrogenase homologs from heterocyst-forming *Nostocales*, which compartmentalize nitrogen fixation^34^. The presence of heterocystous cyanobacteria in the Proterozoic fossil record^32,35^ constraints the minimum ages of Anc3 and Anc4 to approximately 700 Ma to 2.3 Ga (**Figure 1**).

### Functional characteristics of ancestral nitrogenases

The catalytic function of the ancestral nitrogenases was evaluated by their ability to support diazotrophic growth and catalyze acetylene reduction *in vivo* (**Figure 2e-g**) (**Methods**). Relative to the wild-type, all ancestral strains showed lower rates of ethylene production, a standard proxy for N_2_ reduction^36^, reflecting measurable differences in catalytic performance across evolutionary time. (**Figure 2f**).

**Figure 2.**
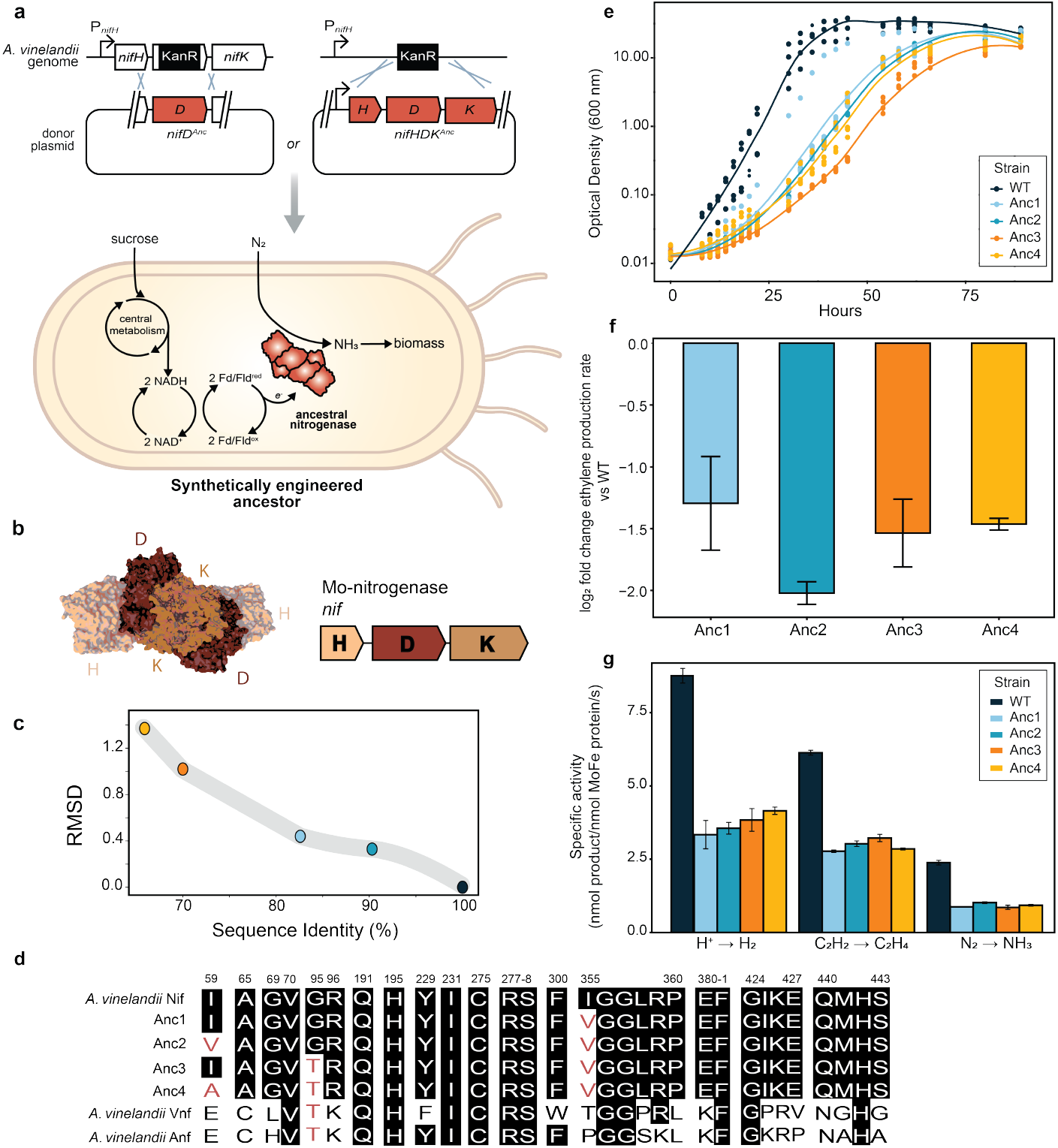
Resurrection and characterization of ancestral nitrogenases. **a**, schematic of methods used to insert ancestral genes into the host *A. vinelandii* genome **b**, Nitrogenase structure and core nif operon. **c**, Root mean square deviation (RMSD) of atoms in predicted structures versus amino acid sequence identity (%). **d**, Sequence alignment of residues within 5 Å of the FeMo-cofactor of NifD. Residue numbering is based on WT *A. vinelandii* NifD. **e**, Growth curves of cultures under nitrogen-fixing conditions. A smoothed curve is shown along with the individual data points of five biological replicates per strain. **f**, *in vivo* ethylene production rates, presented as log2 fold change relative to WT rates. Bars represent mean ± standard error. **g**, *In vitro* specific activity of purified nitrogenases for H^+^, C_2_H_2_, and N_2_ substrates. Bars represent mean ± standard deviation.

Purified ancestral nitrogenases exhibited lower *in vitro* specific activities than their WT counterparts across all tested substrates (**Figure 2g**). Despite this, the engineered strains maintained nitrogen-fixation efficiencies comparable to the WT^37^. Immunodetection showed that nitrogenase protein expressions are generally comparable to the WT **(Extended Data Fig. 3)** and a prior transcriptomic analysis of Anc2 revealed no significant upregulation of nif genes^38^. These results suggest that differences arise primarily from intrinsic functional properties rather than changes in expression.

### Nitrogen Isotope Fractionation: Archean to Now

Batch cultures of *A. vinelandii* strains expressing ancestral nitrogenases were grown under diazotrophic conditions, and cell biomass was harvested for nitrogen isotopic analysis. The isotopic fractionation associated with nitrogen fixation (ε_fixed nitrogen;_ ε^15^N) was calculated relative to atmospheric nitrogen (0‰). All ancestral strains exhibited ε^15^N values ranging from −0.92 to − 2.88‰ (**Figure 3**) and notably all overlap with the range displayed by WT. Anc1 and Anc3 were statistically similar to WT, while Anc2 and Anc4 differed significantly from WT and each other, but there was no discernible trend between fractionation values and increasing ancestral enzyme age (**Figure 3b**). The isotopic fractionations (ε^15^N) associated with the ancestral nitrogenases fall within the extremities of the range reported for modern diazotrophs (+3.7 to −5.86‰) and align with average values observed in extant bacteria (−1.38‰) and archaea (−4.13‰)^16^ (**Figure 4**). Among the variants, Anc3, the second oldest enzyme, exhibited the broadest range in ε^15^N values and the most negative average fractionation.

**Figure 3.**
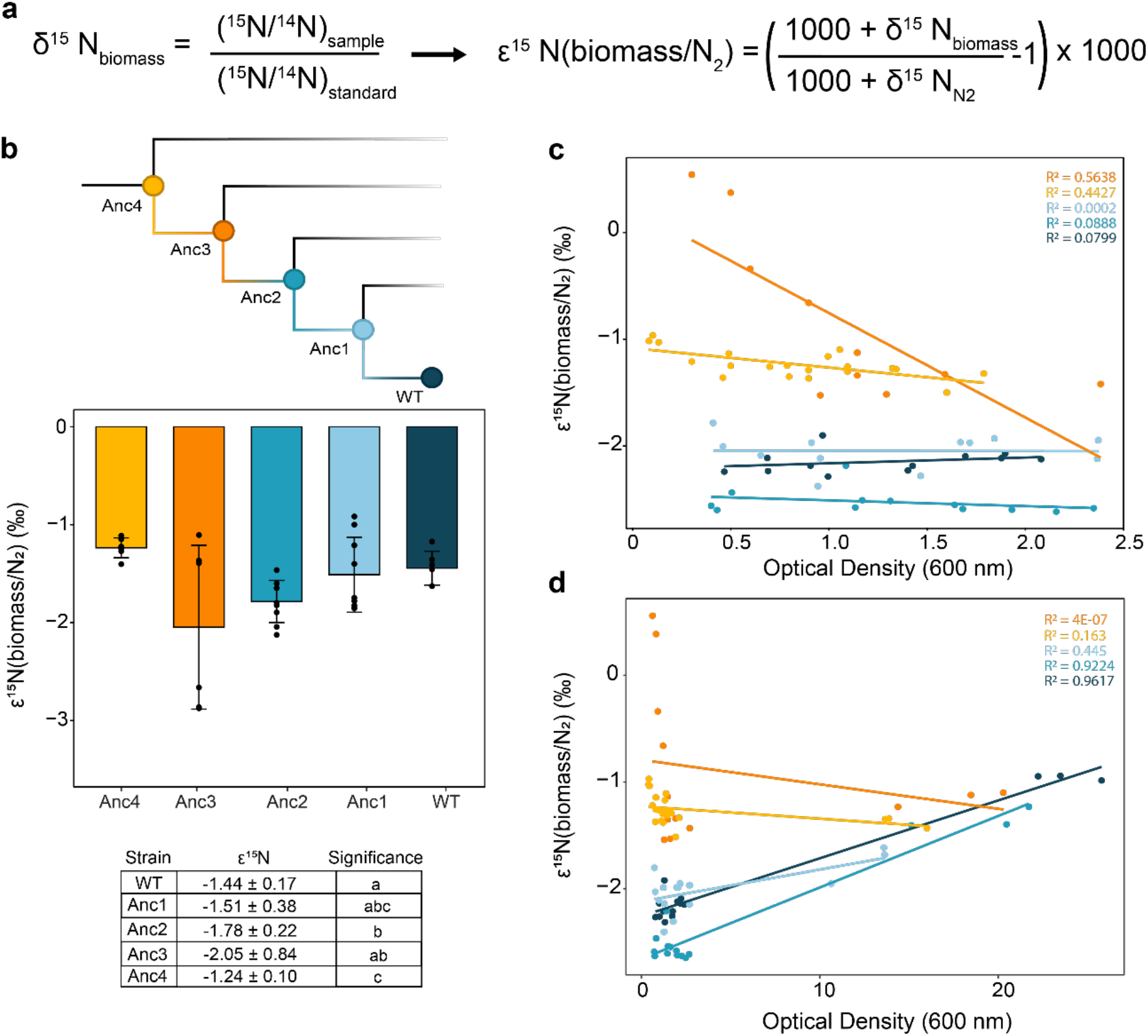
Nitrogen isotope signatures of ancestral nitrogenases. **a**, Equations used to calculate fractionation. **b**, Isotopic fractionation was measured for whole cell biomass of ancestral strains compared to WT *A. vinelandii*. Phylogeny is a subset of Group-I tree modified from Figure 1B. Branches are not to scale. Bars are the average fractionation value calculated across biological replicates, with exact data points shown as well. Error bars represent ±1 standard deviation. Changes in biomass nitrogen isotope signatures throughout early log **c**, and late log to early stationary phase **d**, of diazotrophic growth are shown. The straight lines are the linear regressions.

**Figure 4.**
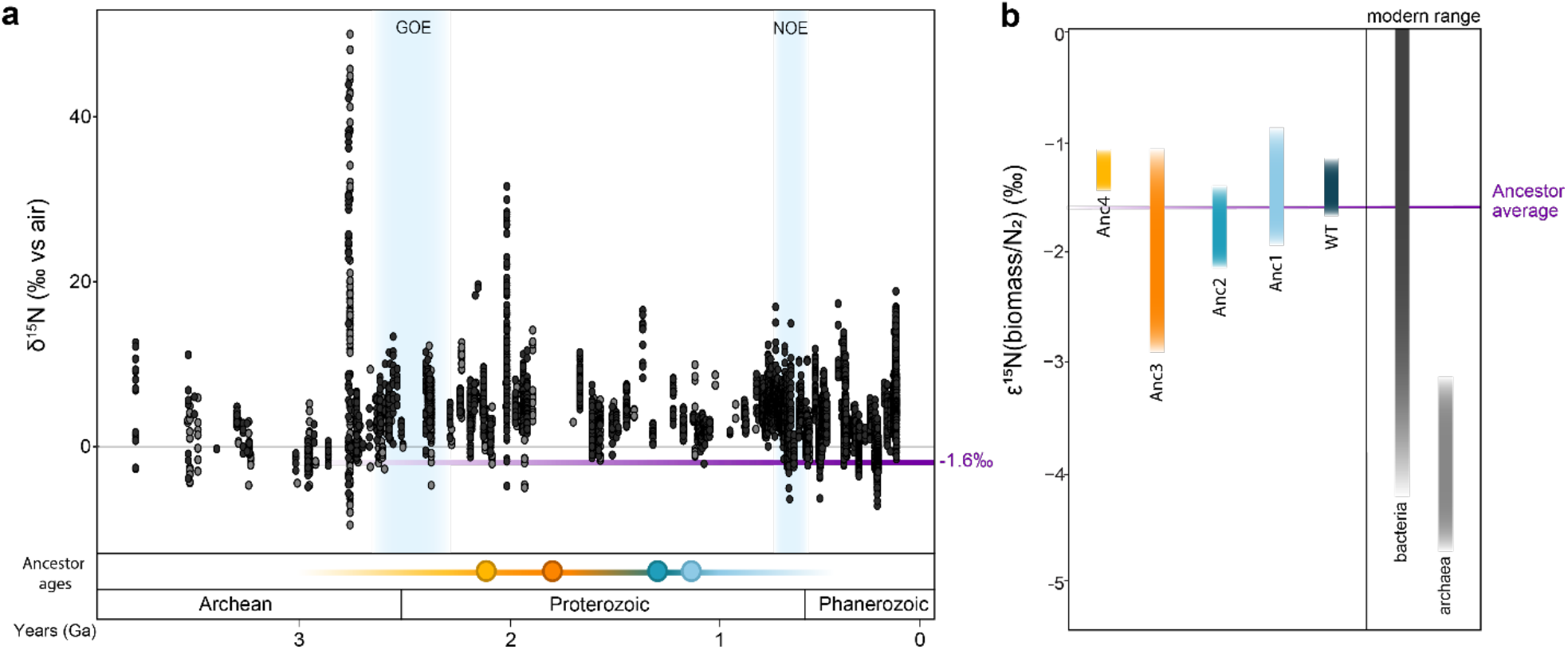
N-isotope fractionation values observed in this study compared to the sedimentary δ^15^N record **a**, and known ranges of extant microbial biomass **b**, The sedimentary database includes bulk rock δ^15^N (dark grey) and kerogen δ^15^N (light grey), from both marine and non-marine rocks across a range of very low to moderate metamorphic grades^22,39,40^. The purple bar in **a** and **b** represents the average fractionation signature of all ancestors in this study and is largely coincident with the minimum d^15^N values seen in the rock record which are presumed to record nitrogen-fixation unaffected by subsequent nitrogen cycling. The partial range of bacterial and archaeal fractionation signatures are labeled as the modern range of N-isotopic values^16^ (see **Extended Data Fig. 5** for full range).

We further assessed whether nitrogen isotopic fractionation changed as a function of diazotrophic growth phase (**Figure 3c-d**). Previous work with WT bacteria has shown up to 2‰ variation in isotopic values throughout growth^14^. In contrast, we observed no significant trend in ε^15^N values throughout early log phase for WT, Anc1, or Anc2, whereas biomass from Anc3 and Anc4 became slightly more ^15^N-depleted (Pearson coefficients = −0.75 and −0.66, respectively). Most strains exhibited <1‰ variation in biomass ε^15^N values, with all strains remaining within the ≤2‰ range reported previously^14^. At late log to stationary phases (i.e., OD_600_ of 8-25), biomass ^15^N increased slightly (i.e., became less ^15^N depleted) for WT, Anc1, and Anc2 (Pearson coefficients = 0.98, 0.67, and 0.96, respectively), while Anc3 and Anc4 showed weak depletion (Pearson coefficients = −0.23 and −0.4, respectively). Overall, isotopic variation across all growth phases was not a primary control on nitrogen isotopic signatures.

## DISCUSSION

In this study, we reconstructed and experimentally characterized the nitrogen isotopic signatures of ancestral nitrogenases to directly probe the molecular origins of Precambrian biosignatures. Because all fixed nitrogen is assimilated into biomass, the measured ^15^N values of cells reflect the isotopic fractionation of the ancestral enzymes. Despite spanning over 2 billion years of evolutionary divergence, the ancestral strains produced biomass ε^15^N values broadly comparable to those of extant nitrogenase. Among them, Anc3 produced the most negative average ε^15^N value, whereas Anc4 - the phylogenetically oldest – produced the least fractionated signal. These results demonstrate that nitrogen isotope fractionation by nitrogenase has remained within a relatively narrow range over deep time, suggesting strong conservation of either enzyme-specific isotope effects or whole-cell regulatory processes. More broadly, this work introduces a new experimental framework for interrogating ancient biogeochemical signatures through synthetic molecular evolution, enabling reconstruction of life’s isotopic fingerprints in deep time.

All measured fractionation values from ancestral strains fall within a narrow range (approximately −1 to −3‰), overlapping both with minimum isotopic signatures observed in Archaean sedimentary rocks (+2 to −2‰^12,41–43^) and with modern Mo-dependent nitrogen fixers (+3.7 to - 5.86‰; bacterial average of −1.38‰^16^) (**Figure 4**). The average across all ancestral strains (−1.6 ± 0.5‰) and the tight clustering of individual strain values suggest a striking isotopic continuity extending deep into Earth’s past. These results support the view that Mo-nitrogenases imparted a consistent isotopic biosignature since at least the Paleoproterozoic, and that they can be distinguished from the larger fractionations associated with V- and Fe-nitrogenases (−6 to − 8‰^13^)^12^.

Critically, whole-rock δ^15^N values capture the integrated signature of ancient microbial communities and ecosystem-scale processing, not just nitrogenase enzymes. Even within diazotrophs, the isotopic fractionation effects of downstream nitrogen assimilation steps remain poorly constrained. Nevertheless, the consistency between ancient enzyme-driven ε^15^N values and the geologic record^22^ (**Figure 4**) suggests that assimilation pathways and intracellular processing steps have also remained broadly conserved over billions of years of evolution. This is comparable to patterns seen in sulfur isotopes, where once-evolved metabolic strategies exhibit long-term evolutionary stasis^9,44,45^.

The similarity of the lowest N-isotope values in early Archean rocks (**Figure 4A**) suggests that the currently accepted minimum age for nitrogenase (3.2 Ga) may be too conservative, and that its origin could predate what is currently resolvable in the rock record. This hypothesis is supported by recent evidence for elevated dissolved Mo concentrations in Archean seawater^46^ and the widespread presence of Mo-dependent enzymes by the mid-Archean^47^. Although Archean sedimentary rocks remain scarce and are often poorly preserved, these constraints do not diminish the significance of our findings. The convergence of geochemical evidence and the consistent ε^15^N signatures inherited from ancestral nitrogenase collectively point to an earlier emergence of Mo-based biological nitrogen fixation than previously considered. It is important to note that while other now-extinct ancient biological strategies for nitrogen fixation cannot be entirely ruled out, there is no evidence for any alternative nitrogenase lineage; all characterized nitrogenases derive from a single ancestral enzyme. Abiotic sources of fixed nitrogen, such as lightning-driven reactions^48^, are also recorded in the rock record, but their isotopic signatures are distinct from those produced by biological nitrogen fixation. Given current knowledge, the most parsimonious interpretation is that the early nitrogen isotope record primarily reflects activity of the nitrogenase enzyme.

Furthermore, while it has been proposed that lower Archean oxygen levels may have enhanced isotopic fractionation via gas diffusion^50^, our data do not support this interpretation. The stability of ancestral nitrogenase-catalyzed fractionation across over 2 billion years when oxygen levels fluctuated considerably suggests that environmental oxygen concentrations did not exert a primary influence on the enzyme’s isotopic discrimination. Instead, our results support an earlier establishment of the biochemical mechanisms underlying nitrogen isotope fractionation, independent of redox conditions. The effects of other environmental factors, such as temperature, on ancestral nitrogenase activity and isotopic fractionation remain to be explored, but the observed biochemical constancy under variable oxygen levels points to a deeply rooted and resilient biochemical mechanism.

These findings point to a broader principle: once a metabolic pathway becomes established and functionally integrated into the biosphere, its core biochemical properties—and the isotopic signatures they produce—can remain remarkably stable over geological timescales. Crucially, this implies that the persistence of a biosignature is governed less by subsequent environmental change and more by the physiological context in which the metabolism originally evolved. Gaining insight into life at this critical point is therefore essential, not only for interpreting ancient isotopic records on Earth, but also for identifying life on other worlds. After all, when life was onto a good thing, it stuck to it.

## ACKNOWLEDGMENTS

This work was supported by the National Aeronautics and Space Administration (NASA) Interdisciplinary Consortium for Astrobiology Research: Metal Utilization and Selection Across Eons, MUSE [80NSSC17K0296] with additional support from the NASA Exobiology Program [NNH23ZDA001N] a part of the NASA LIFE Research Coordination Network, USDA-NIFA Hatch Award [WIS05097], and the UW-Madison Department of Bacteriology Michael and Winona Foster Predoctoral Fellowship award to HR. We would like to thank Amanda Garcia, Scott Chang, Alex Rivier, Chris Wozniak, Kaustubh Amritkar, and Andrew Schauer for their assistance and feedback.

## METHODS

### Ancestral sequence reconstruction

Phylogenetic analysis and ancestral sequence reconstruction of nitrogenase proteins were performed previously^37^. Briefly, nitrogenase protein sequences were collected by BLASTp search^51^ against the NCBI non-redundant protein database (accessed August 2020) and curated manually, yielding 385 sets of H-, D-, and K-subunit sequences. Sequences for each subunit were aligned separately by MAFFT v.7.450^52^. For ancestor Anc2, protein sequence alignments were trimmed using trimAl v.1.2^53^. Evolutionary model selection (LG + G + F) was performed by ModelFinder^54^ implemented in IQ-TREE v.1.6.12^55^. Tree reconstruction (using the trimmed alignment) and ancestral sequence reconstruction (using the untrimmed alignment) were performed by RAxML v.8.2.10^56^ using the best-fit LG + G + F model. For ancestors Anc1, Anc3, and Anc4, an updated tree was built using an untrimmed alignment and RAxML. Ancestral sequence reconstruction was instead performed by PAML v.4.9j^57^ using the LG + G + model. Oxygen metabolism was calculated via the Oxyphen package^58,59^.

### Sequence and structural comparison

The DDKK complex structures for the WT Nif from *A. vinelandii* and the selected Group I ancestors were extracted from the Nitrogenase database (https://nsdb.bact.wisc.edu/)^60^ (**Extended Data Fig. 1)**. To assess the structural variation, we calculated Root Mean Squared Deviation (RMSD) and sequence identity for each ancestor with respect to the WT complex using US-align^61^ (**Figure 2c**).

Sequence embeddings were calculated for the different NifD variant sequences at each node generated during ancestral sequence reconstruction using a 5-dimensional physicochemical representation for each amino acid residue^61^. Dimensionality reduction and clustering visualization were performed using t-distributed stochastic neighbor embedding (t-SNE) from the Scikit-learn implementation^62^ (**Extended Data Fig. 1c**). Mean posterior probability and SH-like aLRT for each residue and node of the ancestral sequences were calculated (**Supplemental Table S3**).

### Strain engineering and functional assessment

Ancestral nitrogenase sequences were obtained from a previously constructed phylogeny^37^. Strains Anc1-3 were constructed in previous studies (Anc1-2^37^, and Anc3^63^). Anc4 was constructed following these previously established methods. Transformants were screened for nitrogen fixation rescue and loss of kanamycin resistance on solid Burk’s (“B”) medium. The growth rate of each strain in nitrogen-fixing conditions was quantified using previously reported methods^64^. In brief, liquid B medium was inoculated from a 24-hour seed culture to an optical density at 600nm of 0.05. Three biological replicates of each strain were tested. The cultures were grown in a 96-well, flat-bottom plate (Greiner Bio-One) with a Breathe-Easy adhesive seal (Diversified Biotech). The plate was incubated at 30°C with a 200-rpm double orbital agitation inside a SPECTROstar Nano Microplate Reader (BMG Labtech, Ortenberg, Germany). An optical density reading at 600nm was taken every 30 minutes for 72 hours. Doubling time and data visualization was conducted using the R package Growthcurver^65^.

Acetylene reduction assay was conducted following previous methods^37^. Biological replicates of each culture were grown overnight and then used to inoculate 100 mL of Burk’s medium with ammonium acetate (“BN media”) to an OD_600_ of 0.01. Cultures were then grown for 16 hours, centrifuged down to a cell pellet at 4700 rpm for 10 minutes, and then resuspended in 100 mL of B medium. The cultures were shaken at 30 °C and agitated at 300 rpm for 4 hours, at which time a rubber septum was placed over each flask. 25 mL of headspace air was removed and replaced with 25 mL of acetylene. Headspace measurements were taken 20, 40, and 60 minutes after acetylene incubation. Ethylene production was quantified via Nexis GC-2030 gas chromatograph (Shimadzu). Cell pellets were collected after the final sampling by centrifuging 24 mL of cultures at 4700 rpm for 10 minutes, washing with 3mL of phosphate buffer, and pelleting once more as above. Cell pellets were stored at –80 °C. The total protein of each pellet was quantified using a Quick Start Bradford Protein Assay kit (Bio-rad) and CLARIOstar Plus plate reader (BMG Labtech). The total protein of each pellet was used to normalize the acetylene reduction rates.

### Growth and sample collection

BN medium was inoculated and grown at 30°C and agitated at 250-300 rpm. Biological replicates (n=6 to 9) were prepared for each strain. After 20-24 hours, the optical density at 600nm (OD_600_) of each culture was measured. These cultures were then used to inoculate 500mL of Burk’s media without any nitrogen (“B media”) in 1L Erlenmeyer flasks to an OD_600_ of 0.01. The cultures were placed in a shaker at 30°C and agitated at 250-300 rpm. The OD_600_ of the cultures was monitored until the cultures reached log or stationary growth.

To collect samples for isotope analysis, 50mL of culture were taken from each flask and centrifuged for 10 minutes at 4700 rpm. The pellet was resuspended and washed with 50mL 1xPO_4_, centrifuging as before. The resulting cell pellets were resuspended and washed again with 1 mL 1xPO_4_ in a benchtop microcentrifuge (5000 rpm for 10 minutes). The supernatant was removed, and the pellets were dried in an oven at 50°C for 48 hours. Laboratory air samples were also collected for analysis.

### Isotopic analyses

All analyses of nitrogen isotopic composition were carried out at the University of Washington Isolab facilities. The bulk nitrogen isotopic analyses were analyzed using an elemental analyzer (Costech ECS 4010) coupled to a continuous-flow isotope-ratio mass spectrometer (CF-IRMS; ThermoFinnigan MAT253).

Approximately 0.500-2.000 mg of freeze-dried *A. vinelandii* biomass powders were weighted into 3.5mm x 5mm tin capsules (Costech) and flash-combusted with 1000°C under an oxygen-rich atmosphere. The resulting N_2_ and CO_2_ gases were purified with chromium oxide at 1000°C to convert trace CO gas to CO_2_, and silvered cobalt oxide at 1000°C to capture sulfur gases and halogens. Then, another purification step was performed with copper at 650°C to convert trace NOx gases to N_2_. Traces of H_2_O were scrubbed with magnesium perchlorate. Subsequently, N_2_ and CO_2_ gases were separated chromatographically in a GC column and analyzed for their isotopic composition.

An aliquot of the lab air (2 mL) was analyzed via a Sercon CryoPrep gas concentration system interfaced to a Sercon 20-20 isotope-ratio mass spectrometer at the University of California, Davis Stable Isotope Facility (**Extended Data Fig. 4**).

Isotope and abundance measurements were calibrated using three in-house standards (two glutamic acids, GA1 and GA2, and dried salmon, SA). In-house standards were calibrated against the international reference materials USGS40 and USGS41^66^. Isotope measurements were reported in standard delta notation relative to Air N_2_:

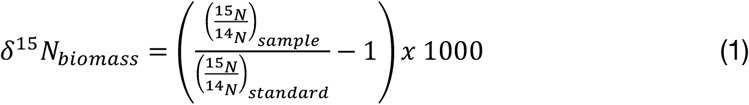

### Calculations and Statistics

ε^15^N of the biomass was calculated using the following equation:

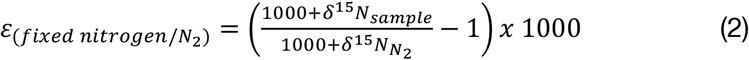

Where δ^15^N_N2_ for the biomass samples is atmospheric N_2_ (0‰).

Statistical significance was calculated using one-way ANOVA, Pearson correlation coefficient, and Tukey’s Honest Significant Difference tests. A significance level of 0.05 was used for all analyses.

## DATA AVAILABILITY STATEMENT

All materials including bacterial strains and plasmids are available to the scientific community upon request. Phylogenetic data, including phylogenetic trees, and ancestral protein sequences are publicly available at https://github.com/kacarlab/N-isotope. All other data are included as extended data and supplementary files.

