## Supplemental Information for "Resurrected nitrogenases recapitulate canonical N-isotope biosignatures over two billion years"

**Table S1.** Molecular resources and strains

| **Reagent type (species) or resource** | **Designation** | **Source or reference** | **Additional information** |
| --- | --- | --- | --- |
| Strain (*A. vinelandii*) | WT | DOI:10.1128/JB.00504–09 | Dennis Dean, Virginia Tech;  Wild-type (WT); Nif^+^ |
| Strain  (*A. vinelandii*) | Δnif | DOI: 10.1128/msystems.00155-24 | Δ*nifHDK*::KanR + Δ*vnfDGK*::StrR + *anfD*::GenR; Nif-, Vnf-, Anf |
| Engineered  Ancestral strain  (*A. vinelandii*) | Anc1 | DOI: 10.7554/eLife.85003 | “Anc1B”; Δ*nifHDK*::*nifHDK*^Anc1B^ (Strep-tagged NifD); Nif^+^ |
| Engineered  Ancestral strain (*A. vinelandii*) | Anc2 | DOI: 10.7554/eLife.85003 | Δ*nifD*::*nifD*^Anc2^ (Strep-tagged NifD); Nif*^+^* |
| Engineered  Ancestral strain (*A. vinelandii*) | Anc3 | DOI:  10.1128/mbio.01271-24 | “AK029”; Δ*nifHDK*::*nifHDKAnc* (Strep-tagged NifD) + Δ*vnfDGK*::StrR + *anfD*::GenR; Nif+, Vnf^-^, Anf^-^ |
| Engineered  Ancestral strain  (*A. vinelandii*) | Anc4 | This study | Δ*nifD*::*nifD*^Anc4^ (Strep-tagged NifD); Nif^+^ |
| Recombinant DNA reagent | pAG19 | DOI: 10.7554/eLife.85003 | *nifHDK*^Anc1B^ (Strep-tagged NifD) + 400-bp *nifHDK* flanking homology regions, synthesized into XbaI/KpnI sites in pUC19; used to construct strain Anc1 |
| Recombinant DNA reagent | pAG14 | DOI: 10.7554/eLife.85003 | *nifD*^Anc2^ (Strep-tagged NifD) +400 bp *nifD* flanking homology regions, synthesized into XbaI/KpnI sites in pUC19; used to construct strain Anc2 |
| Recombinant DNA reagent | pAnc^AK029^ | DOI: 10.1128/mbio.01271-24 | *nifHDK*^Anc3^ (Strep-tagged NifD) + 1000-bp *nifHDK* flanking homology regions, synthesized into XbaI/BamHI sites in pUC19; used to construct strain Anc3 |
| Recombinant DNA reagent | pSC02 | This study | *nifD*^Anc4^ (Strep-tagged NifD) + 1000-bp *nifD* flanking homology regions; synthesized in pUC19; used to construct strain Anc4 |
| PCR primer | 306_nifH_F | DOI: 10.7554/eLife.85003 | GCCGAACGTTCAAGTGGAAA |
| PCR primer | 307_nifH_R | DOI: 10.7554/eLife.85003 | AGAGCCAATCTGCCCTGTC |
| PCR primer | 308_nifD_F | DOI: 10.7554/eLife.85003 | CACCCGTTACCCGCATATGA |
| PCR primer | 309_nifD_R | DOI: 10.7554/eLife.85003 | ACTCATCTGTGAACGGCGTT |
| PCR primer | 310_nifK_F | DOI: 10.7554/eLife.85003 | GCTAACGCCGTTCACAGATG |
| PCR primer | 311_nifK_R | DOI: 10.7554/eLife.85003 | TCAGTTGGCCTTCGTCGTTG |

**Table S2.** Amino acid sequences of reconstructed ancient nitrogenases

| WT NifH | MAMRQCAIYGKGGIGKSTTTQNLVAALAEMGKKVMIVGCDPKADSTRLILHSKAQNTIMEMAAEAGTVEDLELEDVLKAGYGGVKCVESGGPEPGVGCAGRGVITAINFLEEEGAYEDDLDFVFYDVLGDVVCGGFAMPIRENKAQEIYIVCSGEMMAMYAANNISKGIVKYANSGSVRLGGLICNSRNTDREDELIIALANKLGTQMIHFVPRDNVVQRAEIRRMTVIEYDPKAKQADEYRALARKVVDNKLLVIPNPITMDELEELLMEFGIMEVEDESIVGKTAEEV* |
| --- | --- |
| WT NifD | MTGMSREEVESLIQEVLEVYPEKARKDRNKHLAVNDPAVTQSKKCIISNKKSQPGLMTIRGCAYAGSKGVVWGPIKDMIHISHGPVGCGQYSRAGRRNYYIGTTGVNAFVTMNFTSDFQEKDIVFGGDKKLAKLIDEVETLFPLNKGISVQSECPIGLIGDDIESVSKVKGAELSKTIVPVRCEGFRGVSQSLGHHIANDAVRDWVLGKRDEDTTFASTPYDVAIIGDYNIGGDAWSSRILLEEMGLRCVAQWSGDGSISEIELTPKVKLNLVHCYRSMNYISRHMEEKYGIPWMEYNFFGPTKTIESLRAIAAKFDESIQKKCEEVIAKYKPEWEAVVAKYRPRLEGKRVMLYIGGLRPRHVIGAYEDLGMEVVGTGYEFAHNDDYDRTMKEMGDSTLLYDDVTGYEFEEFVKRIKPDLIGSGIKEKFIFQKMGIPFRQMHSWDYSGPYHGFDGFAIFARDMDMTLNNPCWKKLQAPWEASEGAEKVAASA* |
| WT NifK | EPASR*NQSQLPAVPRSGLQGHACQEARRLRGKVSAGQDRRSIPVDHHQGIPGAELPARSPDRQPGQGLPAAGRRSLRPRFREDHALRARFPGLRRLLPLLLQPSFPRAGFLRFRLHDRRRGSVRRPAEHEGRSAEL*GYLQARHDRSVHHLHGRGHR*RPQRLHQQLEEGRFHS*RVPGAVRPYPELRGQPRDRLGQHVRRHCSLLHPEVHGRQGGWQQQEDQHRPRLRDLPGQLPRDQAHAFGNGRGLQPALRSGRSAGHPG*RPVPHVRGRHHSGRDEGRSERPQHRPAAAVAPGEDQEVRRGYLEARSTEAEHPDGPGLDRRVPDESQRNQRPADSGEPDQGAWPSGRHDDRLPHLAARQAFRPVG*SGLRDGPGQVPAGTGLRAGTHSLPQRQQALEEGGRRHPRRFALRQECYRLHRQGPVAPAFAGLHRQAGLHDRQQLR*VHPARHPAQGQGVRGSADPYRLPDLRPSSPASLHHPGLRGRHADPDHPGELDPGTSGRGNPRYAGHRLQPRPGTL* |
| Anc1 NifH | MAMRQCAIYGKGGIGKSTTTQNLVAALAEAGKKVMIVGCDPKADSTRLILHSKAQNTIMEMAAEAGTVEDLELEDVLKAGYGDIKCVESGGPEPGVGCAGRGVITAINFLEEEGAYEDDLDFVFYDVLGDVVCGGFAMPIRENKAQEIYIVCSGEMMAMYAANNISKGIVKYANSGGVRLAGLICNSRNTDREDELIMALAEKLGTQMIHFVPRDNVVQRAEIRRMTVIEYDPKAKQADEYRALAQKIIDNKMLVIPTPITMDELEELLMEFGIMDEEDESIVGKTAAEL* |
| Anc1 NifD | MSAASWSHPQFEKMSRDEVEALIQEVLEVYPEKAKKDRAKHLAVNDQSVEQSKKCITSNKKSLPGVMTIRGCAYAGSKGVVWGPIKDMIHISHGPVGCGQYSRAGRRNYYIGTTGVNAFVTMNFTSDFQEKDIVFGGDKKLAKLIDEIETLFPLNKGISVQSECPIGLIGDDIEAVAKQKSAELGKTVVPVRCEGFRGVSQSLGHHIANDAVRDWVLSKRDDDDSFESTPYDVAIIGDYNIGGDAWSSRILLEEMGLRVVAQWSGDGTISEMELTPKVKLNLVHCYRSMNYISRHMEEKYGIPWMEYNFFGPTKTIESLRKIAAQFDESIQAKCEEVIAKYKPEWEAVIAKYRPRLEGKRVMLYVGGLRPRHVIGAYEDLGMEVVGTGYEFAHNDDYDRTIKEMGNATLLYDDVTGYEFEEFVKKVKPDLIGSGIKEKYIFQKMGIPFRQMHSWDYSGPYHGFDGFAIFARDMDMTLNNPCWKKLQAPWKKAESEAEAVAASA* |
| Anc1 NifK | EPASR*HQTELPAVPR*GVQGHACQEARQLRGKAPAGEDRRSIPVDHHRGIPGAELPARSPDRQPGQGLPAAGRRSLRPRFREDHALRARFPGLRRLLPLLLQPSFQGADLLRFRLHDRRRGSVRRPAEHEGRSGEL*GYLQARHDRSVHHLHGRGHR*RPQRLHQQLEEGRSHS*GVPGAVRPYPELRGQPHHRLGQHVRRHCSLLHPELHGGQGGWQQRQDQHRPRLRDLPGQLPRDQAHDERNGRGLHPALRSGRSAGHPG*RPVPHVRGRHHSGRDQGRSERPQHPAAAAVAADQDQEVRQEHLEARSPEAEHPDGPGVDRRVPDESQRNHRQADSGEPGQGAWPSGRHDDRLPHLAARQEVRPVG*SGLRDGHDQVPAGTGLRADPHSLQQRQQALEEGDGGHPRRVALRRQ*RGPHRQGPVAHAFAGLHQQAGLHDRQQLR*VHPARHPVQGQGVRGSADPYRLPDLRPSSPASPDHPGLRGRHADPDHPGELGAGTSGRGNPRYAGHRLQLRPGTL* |
| Anc2 NifD | MSTASWSHPQFEKMTREETQALIQEVLEVYPEKARKDRAKHLAVNDPSIEQSKKCITSNRKSLPGVMTVRGCAYAGSKGVVWGPIKDMIHISHGPVGCGQYSRAGRRNYYTGTTGVNTFGTMNFTSDFQEKDIVFGGDKKLAKIIDEIETLFPLNKGISVQSECPIGLIGDDIEAVAKKASKEIGKPVVPVRCEGFRGVSQSLGHHIANDAIRDWVLDKRDGQSFESTPYDVAIIGDYNIGGDAWSSRILLEEMGLRVVAQWSGDGTLAEMENTPKVKLNLLHCYRSMNYISRHMEEKYGIPWMEYNFFGPTKIAESLRKIAAHFDDTIQENAERVIAKYQPMMEAVIAKYRPRLEGKKVMLYVGGLRPRHVIGAYEDLGMEVVGTGYEFAHNDDYDRTIKELGDATLLYDDVTGYELEEFVKRLKPDLIGSGIKEKYIFQKMGIPFRQMHSWDYSGPYHGYDGFAIFARDMDMTLNNPCWNKLTPPWKKT* |
| Anc3 NifH | MALRQIAFYGKGGIGKSTTSQNTVAALAEMGKKVMIVGCDPKADSTRLILHAKAQTTVMQLAAEAGSVEDLELEDVLKTGYGGIKCVESGGPEPGVGCAGRGVITAINFLEEEGAYEEDLDFVSYDVLGDVVCGGFAMPIREGKAQEIYIVTSGEMMAMYAANNISKGILKYANSGGVRLGGLICNSRNTDREDELIEALAKRLGTQMIHFVPRDNVVQHAELRRMTVIEYSPESKQADEYRTLAKKIIDNKMLVIPTPITMDELEDLLMEFGIMEEEDESIVGKTAAEA* |
| Anc3 NifD | MSTASWSHPQFEKKTTKEQTQELIEEVLEAYPEKARKDRAKHLAVNDPEEEGSSSCSVKSNIKSRPGVMTIRGCAYAGSKGVVWGPIKDMIHISHGPVGCGQYSWGTRRNYYNGTTGVDNFVTMQFTSDFQEKDIVFGGDKKLAKIIDEIEELFPLNKGISIQSECPIGLIGDDIEAVAKKKSKEIGKPVVPVRCEGFRGVSQSLGHHIANDAIRDWVLDKRDKEFEPTPYDVAIIGDYNIGGDAWSSRILLEEMGLRVIAQWSGDGTLNEMANTPKAKLNLIHCYRSMNYICRHMEEKYGIPWMEYNFFGPTKIAESLRKIAAHFDEKIQENAEKVIAKYQPQMDAVIEKYRPRLEGKKVMLYVGGLRPRHVIGAYEDLGMEVIGTGYEFAHNDDYQRTTEELKDGTLIYDDVTAYELEEFVKKLKPDLVGSGIKEKYVFQKMGIPFRQMHSWDYSGPYHGYDGFAIFARDMDMAINSPVWNLIKAPWKKA* |
| Anc3 NifK | EPERRENQRPQRAVPAGGVPGDVREQAQAVRERPQRRGGGGSSRVDQDRGIQGEELRPRSPDHQPGQGLPAAGRRSRRPRFRGHPALRARFPGLRRLLPLPLQPSFQGAVPGRFQLHDRRRGSVRRPEQHDRGSAERLRSVQAQDDRSVHHLHGRGHR*RPQRLHQERQGGR*HSSRLPGAVRPYPELRGQPHHRLRQHDEGHSEPPDRGQGGRDHQRQDQHHPRLRHLHRQLPRDQAHP*PDGRGLHHPRRSQRRAGQPG*RRVQDVPGRHHSGRGEGRYQRQGHHQPAEVQHHQDRRVHQEEVEAADRGREHPDRHQGHRRVPDENQRTDRQADSGGAGEGAWPSGRRHDRLPPVDSRQEVRHLR*SGPGAGPDQLPAGNGCRAGTHSLHQRQQEVGEGDAGPARQLALRPGGSGLGRQGPVAPAFAAVHRAGGLPDRQQLR*VPGPRHRHPADPYRLPDLRPSSPASLPDHRLPGRPEPADLDREHHPGRAGPQHQEDRLQLRPGTL* |
| Anc4 NifD | MSTASWSHPQFEKKEITKEQTQKLIEEVLEVYPEKARKDRAKHLAVNDPESSCAVKSNVKSRPGVMTARGCAYAGSKGVVWGPIKDMIHISHGPVGCGHYSWGTRRNYANGTTGVDNFVTFQFTSDFQEKDIVYGGDKKLEQICREIKELFPLAKGISIQSECPVGLIGDDIEAVAKKMSKELGIPVVPVRCEGFRGVSQSLGHHIANDAIRDHVLGKRELEFEPTPYDVAIIGDYNIGGDAWASRKILEEMGLRVIAQWTGDGTINELATTHKAKLNLIHCYRSMNYICKHMEEKYGIPWMEYNFFGPTKIYESLRKIAAHFDDKIQENAEKVIAKYQPMMDAVIEKYRPRLEGKKVMLYVGGLRPRHVIGAYEDLGMEVIGTGYEFAHKDDYERTYPEMKEGTLIYDDVTEYELEEFVKKLKPDLVGSGIKEKYVFQKMGIPFRQMHSWDYSGPYHGFDGFPIFARDMDMAINSPTWNLIKAPWKKE* |

**Table S3**. Mean posterior probabilities and branch support values for reconstructed ancestral nodes

| **Ancestor** | **Mean posterior probability** | **SH-like aLRT** |
| --- | --- | --- |
| Anc1 | 0.984 | 99 |
| Anc2 | 0.983 | 99 |
| Anc3 | 0.960 | 100 |
| Anc4 | 0.939 | 86 |

**Table S4**. Internal standards used in EA-IRMS analysis

|  | **Glutamic Acid (GA1)** | **Glutamic Acid (GA2)** | **Bristol Bay Sockeye Salmon** |
| --- | --- | --- | --- |
| **Run** | Corrected δ^15^N (permil) ± standard deviation | Corrected δ^15^N (permil) ± standard deviation | Corrected δ^15^N(permil) ± standard deviation |
| 1 | -4.6 ± 0.093726 | -5.7 ± 0.078051 | 11.3 ± 0.050628 |
| 2 | -4.4 ± 0.49298 | -5.7 ± 0.24341 | 11.3 ± 0.10804 |
| 3 | -5.3 ± 0.33768 | -5.7 ± 0.073993 | 11.3 ± 0.07077 |
| 4 | -4.5 ± 0.093628 | -5.7 ± 0.097292 | 11.3 ± 0.10299 |
| 5 | -4.7 ± 0.23872 | -5.7 ± 0.13916 | 11.3 ± 0.18916 |
| 6 | -4.6 ± 0.17467 | -5.7 ± 0.18533 | 11.3 ± 0.19046 |
| 7 | -4.7 ± 0.13963 | -5.7 ± 0.27915 | 11.3 ± 0.03863 |
| 8 | -4.5 ± 0.16433 | -5.7 ± 0.074317 | 11.3 ± 0.10502 |
| 9 | -4.6 ± 0.092177 | -5.7 ± 0.16752 | 11.3 ± 0.16707 |

**Table S5**. Accuracy and precision of EA-IRMS data based on internal standards

| **Run** | **Property** | **Accuracy** | **Precision** |
| --- | --- | --- | --- |
| 1 | δ^15^N (permil) | 0.0029448 | 0.093726 |
|  | % N | -0.11458 | 0.10448 |
| 2 | δ^15^N (permil) | 0.21543 | 0.49298 |
|  | % N | -0.81937 | 0.87955 |
| 3 | δ^15^N (permil) | -0.73393 | 0.33768 |
|  | % N | -0.55245 | 0.11179 |
| 4 | δ^15^N (permil) | 0.10494 | 0.093628 |
|  | % N | -0.12556 | 0.13197 |
| 5 | δ^15^N (permil) | -0.076057 | 0.23872 |
|  | % N | -0.3233 | 0.068468 |
| 6 | δ^15^N (permil) | -0.042714 | 0.17467 |
|  | % N | -0.087611 | 0.056225 |
| 7 | δ^15^N (permil) | -0.10577 | 0.13963 |
|  | % N | -0.67498 | 0.12777 |
| 8 | δ^15^N (permil) | 0.053788 | 0.16433 |
|  | % N | -0.22142 | 0.094699 |
| 9 | δ^15^N (permil) | -0.0079878 | 0.092177 |
|  | % N | -0.062411 | 0.091385 |

**Table S6.** Fractionation values for strain biomass analyzed via EA-IRMS

| Strain | ε^15^N vs Air N_2_ (permil) |
| --- | --- |
| WT | -1.1744 |
|  | -1.6292 |
|  | -1.4107 |
|  | -1.6299 |
|  | -1.359 |
|  | -1.4582 |
| Anc1 | -1.7816 |
|  | -1.8266 |
|  | -1.4022 |
|  | -1.8561 |
|  | -1.8512 |
|  | -1.7486 |
|  | -1.2126 |
|  | -0.91809 |
|  | -1.002 |
| Anc2 | -1.4655 |
|  | -1.6016 |
|  | -1.8978 |
|  | -1.6498 |
|  | -1.7995 |
|  | -1.6482 |
|  | -2.0456 |
|  | -2.1271 |
|  | -1.825 |
| Anc3 | -1.3686 |
|  | -1.3942 |
|  | -1.1081 |
|  | -2.8656 |
|  | -2.8771 |
|  | -2.6646 |
| Anc4 | -1.1146 |
|  | -1.1502 |
|  | -1.2261 |
|  | -1.2465 |
|  | -1.4052 |
|  | -1.2733 |
